# Identification of novel differentially expressed zooxanthellal genes from *Aiptasia-Symbiodinium* endosymbiosis through SDS-based RNA purification

**DOI:** 10.1101/2020.09.01.277251

**Authors:** Bo-Nien Chen, Paching Song, Ming-Chyuan Chen, Ming-Chang Hong

## Abstract

Endosymbiosis between dinoflagellates and cnidarian hosts first occurred more than 200 million years ago; however, symbiosis-specific genes and cellular processes involved in the establishment, maintenance, and breakdown of endosymbiosis remain unclear. Therefore, this study aimed to identify the zooxanthellal genes associated with the aforementioned biological processes during endosymbiosis in *Aiptasia-Symbiodinium* endosymbionts. Here, zooxanthellae isolates were treated with 0.02% SDS to decrease potential host RNA contamination and to enhance the identification of novel symbiosis/nonsymbiosis-associated differentially expressed zooxanthellal genes through suppressive subtractive hybridization (SSH) and next-generation sequencing (NGS) methods. Consequently, among 214 symbiosis-specific transcripts identified herein that displayed identity to only 5.6% of host-derived transcripts, 64% were well-known functional genes. In the nonsymbiotic stage, 181 differentially expressed transcripts were identified, of which 64.1% belonged to well-known functional genes. BLAST revealed that 8 categories of cellular processes were significantly induced in symbiotic or nonsymbiotic zooxanthellae. Together with the results of quantitative analysis, the results revealed that photosynthesis, flagellate biosynthesis and motility, stress-induced responses, cell wall biosynthesis, starch synthesis and transport, lipid biosynthesis and metabolism, host/symbiont immune response, intercellular communication, cell growth, and cell cycle regulation were the major cellular processes occurring in symbiotic/nonsymbiotic stages. The present results provide insights into the mechanisms involved in regulating the different physiological processes in symbiotic/nonsymbiotic zooxanthellae and may guide future studies.

## Introduction

Approximately one-third of all marine species reside in coral reef ecosystems and contribute to one-fourth of the produce of marine fisheries. The maintenance of coral reef ecosystems is primarily based on the association between corals and symbiotic photosynthetic dinoflagellates (*Symbiodinium* spp.). The single-cell symbiotic dinoflagellate is concealed within the organelles of the gastric dermis tissues within the host coral cells, called symbiosomes [1,2]. Symbiotic associations have numerous benefits. From symbiotic bodies, corals receive amino acids, fatty acids, and photosynthetic products including glucose, and symbionts effectively utilize nitrogen- and phosphorus-containing waste from host cells [3–7]. The maximal calcification rate of coral reefs has resulted from endosymbiosis [8–10]. Furthermore, symbiotic dinoflagellates prevent harmful ultraviolet radiation from damaging corals through mycin-like amino acids (MAAs) [11,12]. However, endosymbiosis is often disrupted owing to environmental stresses including high temperature, high CO_2_ concentration, and pollution [13–16], resulting in coral bleaching [13,17]. Coral bleaching represents breakdown of the symbiotic integrity between corals, zooxanthellae, and potential microbial communities, leading to negative effects on coral health [18]. However, details regarding numerous interactions including the intracellular and molecular responses during symbiosis establishment, maintenance, and disintegration remain unclear.

Currently, successfully surviving zooxanthellae have to face the complicated host symbiosome microenvironment, which provides resources for growth, sustenance, and reproduction [19]. Previous studies have reported the correlation between photosynthesis and carbonic anhydrase activity, indicating that sources of inorganic carbon required for zooxanthellae are actively provided by the coelenterate host [20,21]. Symbiotic dinoflagellates use phosphatase to hydrolyse the phosphate esters of the inorganic phosphates of the coelenterate host and then transport them into cells through transport proteins [22–25]. Regarding the essential nitrogen source for plants, studies have reported that ammonia waste produced by the host is transferred to the symbiosomes and then converted to ammonium ions in the acidic lumen, similar to the highly acidic inner cavity of the symbiosome, which is markedly different from seawater [26,27], or it might be transported through transport proteins into symbiosomes to facilitate the use of zooxanthellae [24,26,28]; however, the underlying mechanism is unclear. Furthermore, previous studies have reported that the TCN-β signalling pathway and Tsr protein are involved in the maintenance of coelenterate hosts and symbiotic tissues [29,30]. Furthermore, the cell cycle of symbiotic dinoflagellates is arrested in the G_1_ phase for a longer period than that in the nonsymbiotic form and is not regulated by host-derived inorganic nutrients [31,32]. These associations indicate potential physiological responses and pathways of symbiotic dinoflagellates during endosymbiosis.

Recent studies have reported several potential symbiosis-related genes. Coral lectin binds to specific cell wall glycans on zooxanthellae, and their interactions serve as a potential key mechanism for the host to obtain specific symbionts [33,34]. Corals are coated with symbiotic corpuscular membranes of symbiotic algae. Membrane-associated RAB family proteins help retain healthy zooxanthellae cells in the symbiosomes and exclude dysfunctional symbionts [35–37]. Furthermore, the genetic and molecular mechanisms in cnidarian and dinoflagellate endosymbionts have received increasing attention [38]. Next-generation sequencing (NGS) technology has revealed genomic sequences of two nonsymbiotic cnidarians, *Nematostella vectensis* and *Hydra magnipapillata* [39,40], and two symbiotic algae, *Symbiodinium kawagutii* and *S. minutum* [41,42]. Numerous studies have performed transcriptome analysis for coelenterates including *Acropora millepora, Acropora palmata, Pocillopora damicornis*, and *Aiptasia pallida* [43–47]. These data have helped understand the interactions among genes. However, observations regarding symbiosis-specific gene responses between symbiotic and nonsymbiotic states seem slightly different [48,49], potentially indicating that pure genomic analyses have not effectively revealed symbiosis-associated genes and potential cellular responses differing between the symbiotic and nonsymbiotic states. Hence, further improvement of the method is needed to obtain a molecular response truly reflecting endosymbiosis.

NGS can provide large-scale transcriptome data on various types of corals or their symbiotic algae. However, handling symbiotic zooxanthellae in host cells is difficult owing to the possibility of host RNA contamination during nucleic acid extraction. The presence of host RNA can interfere with sequencing reactions and reduce accuracy and miscalculate minor but important symbiotic gene abundances. This study aimed to identify the zooxanthellal genes associated with the aforementioned biological processes during endosymbiosis in *Aiptasia-Symbiodinium* endosymbionts. In this study, we treated *Aiptasia-Symbiodinium* endosymbionts with SDS before nucleic acid extraction to reduce the amount of host RNA contamination and performed combinatorial suppressive subtractive hybridization (SSH)-NGS sequencing to more accurately identify potential symbiotic and nonsymbiotic gene groups. The potential molecular responses provide more accurate information for establishing and maintaining intracellular endosymbionts.

## Materials and methods

### Animals and dinoflagellates

Cnidarian *A. pulchella* and its symbiont, *Symbiodinium spp.* Clade B, were obtained from a private aquaculture farm in Pingtung and maintained in an aquarium of 160 × 60 × 75 cm^3^ with 470 L of water. Salinity, temperature, and pH were maintained at 35 psu, 26–28 °C, and pH 8.0–8.5, respectively, with access to natural sunlight. Frozen shrimp was fed once a week as bait. Each month, half the volume of water was replaced with fresh sea water. To induce bleaching in the sea anemones, symbiotic sea anemones were treated with cold shock at 4 °C for 1 h and maintained in a dark aquarium until bleaching occurred. No residual zooxanthellae were detected with the bleaching sea anemones upon microscopic analysis. Nonsymbiotic zooxanthellae were harvested from symbiotic sea anemones through homogenization, centrifugation, and resuspension. After zooxanthellae were purified, the free-living zooxanthellae were cultured in a T75 flask containing f/2 medium with additional 100, 50, and 50 μg/ml ampicillin, kanamycin, and streptomycin. Zooxanthellae were cultured at 26 °C and on a 12/12-h light/dark cycle.

### RNA purification and cDNA preparation

Total RNA of *A. pulchella* and *Symbiodinium spp.* were extracted using TriZol (Thermo Fisher Scientific, USA). Additionally, before RNA extraction from *Symbiodinium spp.*, SDS treatment was performed for 10 seconds. In total, 5 μg of total RNA of both *A. pulchella* and *Symbiodinium spp.* was used to synthesize ss cDNAs using SMARTer™ PCR cDNA Synthesis Kit (Takara Bio, USA). Furthermore, smart-IV primers were used: (5’-AAGCAGTGGTATCAACGCAGAGTGGCCATTACGGCCGGG-3’) and CDSIII primers [5’-ATTCTAGAGGCCGAGGCCCCCGGGGGGCC-d(T)30N–1N-3’] for reverse transcription and synthesis of ds cDNA gene libraries.

### SSH analysis

Before SSH, both symbiotic and nonsymbiotic zooxanthellae were cultured in seawater at 26 °C at a salinity of 35 psu and pH 8.0–8.5 with a light intensity of 80–100 μmol photons m^−2^s^−1^ for 2 weeks to prevent environmental differences. After acclimatization, RNA extraction and cDNA construction were performed and 4 genes including β-actin of *A. pulchella,* and β-actin, PCP, and HSP90 of *Symbiodinium spp.* were assessed through semi-quantitative PCR analysis to evaluate potential environmental stress-induced responses. Thereafter, SSH was performed using a PCR-Select™ cDNA Subtraction Kit (Takara Bio). The differentially expressed genes thus determined were then cloned into the pCR®II-TOPO vector and transformed into Mach1™-T1R competent cells for sequencing. The sequences thus obtained were extended with a home-curated NGS-based gene database of *A. pulchella* and *Symbiodinium spp.* clade B and then were aligned with NCBI GenBank database, using BLASTX [50]. Furthermore, multiple sequence alignment and phylogenetic tree analysis were performed using DNASTAR software [51].

### NGS analysis

Non–co-symbiotic symbiotic algae and albino beautiful anemones were selected for RNA extraction. Thereafter, 30-min DNase I treatment was administered at 37 °C, followed by RNA purification using TriZol. Thereafter, a Chroma spin+TE-400 column was used to eliminate small RNA. Finally, after electrophoresis with 1% agarin gum, QC analysis was outsourced, and NGS analysis was performed by BGI Genomic company (China).

### Real-time quantitative PCR

The primer annealing temperature range for each gene was set at 57–63 °C, the expected product size was 50–150 bp, the final primer concentration in the reaction mix was 100 nM, and the mRNA content was 5 ng. The detail of primer sequences is all provided in supplementary information (Table S3). The real-time PCR reagent was 2× SYBR green master mix (Toyobo, Japan), and the total reaction volume was 25 μL. The cycling conditions were as follows: pre-denaturation at 95 °C for 1 min, denaturation at 95 °C for 15 s, and extension at 60 °C for 1 min, followed by melting curve analysis. The machine model used herein was the SYBR 48-well template StepOneTM system (ABi), and the analysis software was StepOne software v2.1.

## Results and discussion

### Host RNA contaminations was effectively reduced through SDS treatment

To reduce host RNA contamination, which may affect the accuracy and accuracy of the deduced hybridization, we compared the differentially expressed genes between symbiotic and nonsymbiotic zooxanthellae. After symbiotic/nonsymbiotic zooxanthellae were cultured and acclimated under the same conditions, RNA extraction was performed. During the extraction of symbiotic zooxanthellae, the homogenate of the symbiotic sea anemone was treated with 0%, 0.01%, and 0.02% SDS for 30 s, followed by coupling with suspension and centrifugation until the residual host cell and tissue debris were effectively eliminated. Semi-quantitative reverse transcription PCR was performed to detect host RNA contamination levels. Consequently, 0.02% SDS treatment significantly decreased *Aiptasia* actin signalling but did not affect zooxanthellal actin, HSP90, and PCP expression (Fig 1), indicating that SDS neither causes marked stress to the zooxanthellae nor induces differential gene expression. These results indicate an effective reduction in anemone host RNA contamination, which is conducive for SSH. These results indicate that SDS effectively binds lipid molecules; thus, nucleic acids (DNAs and RNAs) become exposed in cells with damaged membranes; these nucleic acids are then exposed to the aqueous homogenate fluid and digested by endogenous RNases and DNases [52]. However, the zooxanthellal DNAs and RNAs are potentially protected by their cell wall and cannot be significantly damaged during short periods. Therefore, SDS treatment would benefit the purification of the zooxanthellae population.

**Fig 1.**
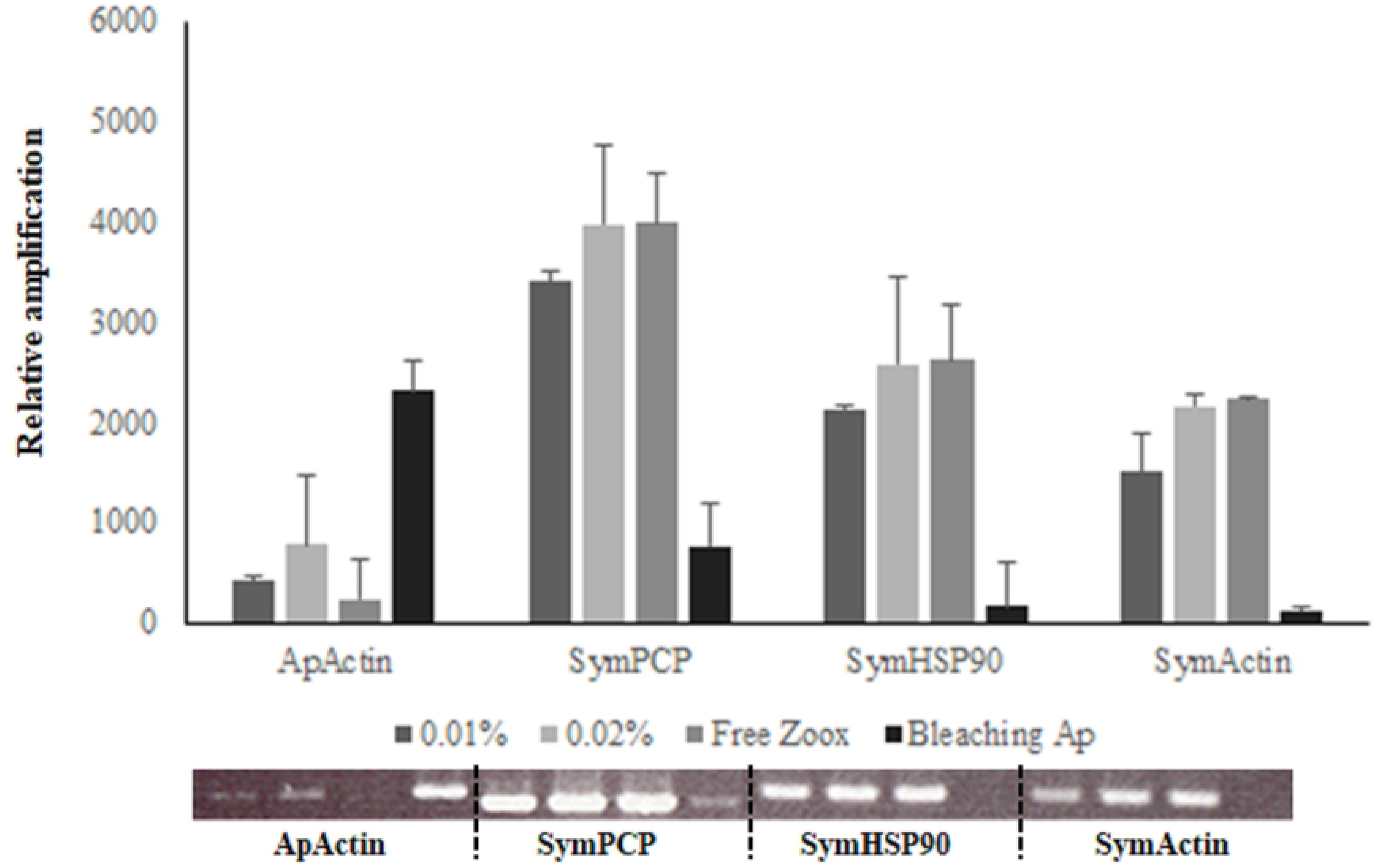
SDS treatment-based RNA purification in zooxanthellae. When harvesting hosts harbouring zooxanthellae, 0.01% and 0.02 % SDS treatment were administered during RNA isolation to efficiently eliminate host RNA contamination. Thereafter, primers specific for actin, PCP, and HSP90 were assessed through semi-quantitative RT-PCR analysis to reveal potential host RNA contamination. These results indicate that 0.02% SDS treatment efficiently decreased host RNA contamination without significant stress on other intracellular molecular responses. ‘ApActin’ represents *Aiptasia pulchella* actin; ‘SymPCP, SymHSP90, and SymActin’ represent zooxanthellal chloroplast soluble peridinin-chlorophyll a-binding protein precursor, heat shock protein 90, and actin, respectively. ‘Free Zoox’ indicate the cDNA derived from cultured nonsymbiotic zooxanthellae, and ‘Bleaching Ap’ was derived from bleaching *Aiptasia pulchella*.

### Transcriptome assay of bleaching *A. pulchella* revealed a 5.6% reduction in host-derived SSH-harvested sequences

SSH is a potent method for analysing differences in gene expression patterns between symbiotic and nonsymbiotic zooxanthellae. Herein, SSH was coupled with an NGS-based transcriptome assay and SDS treatment to further expand the zooxanthellae gene database and effectively reduce host RNA contamination to better identify novel and unknown genetic and cellular responses. As shown in Fig 2, to confirm the reduction in host RNA contamination, the database of the bleaching anemone and nonsymbiotic zooxanthellae gene pool curated through NGS allowed for the extrapolation and identification of DNA sequences generated through SSH. After sequencing was completed, 69,858 conserved sequences of bleaching *Aiptasia* anemones were obtained, including 17,707 contigs and 52,151 unigenes; and 69,327 conserved sequences of isolated free-living *Symbiodinium* zooxanthellae, including 6,434 contigs and 62,893 unigenes, respectively. Details regarding Clusters of Orthologous Genes (COG) are provided in Figs 3 and 4. In total, 214 differentially expressed genes were detected in the symbiotic stage and 181 genes were detected in the nonsymbiotic stage. Only 12 genes (5.6%) in the symbiotic stage expressed zooxanthellal genes and belonged to *A. pulchella* (Fig 5). Altogether, our results indicate that 0.02% SDS treatment helps effectively reduce host RNA contamination and their potential interference.

**Fig 2.**
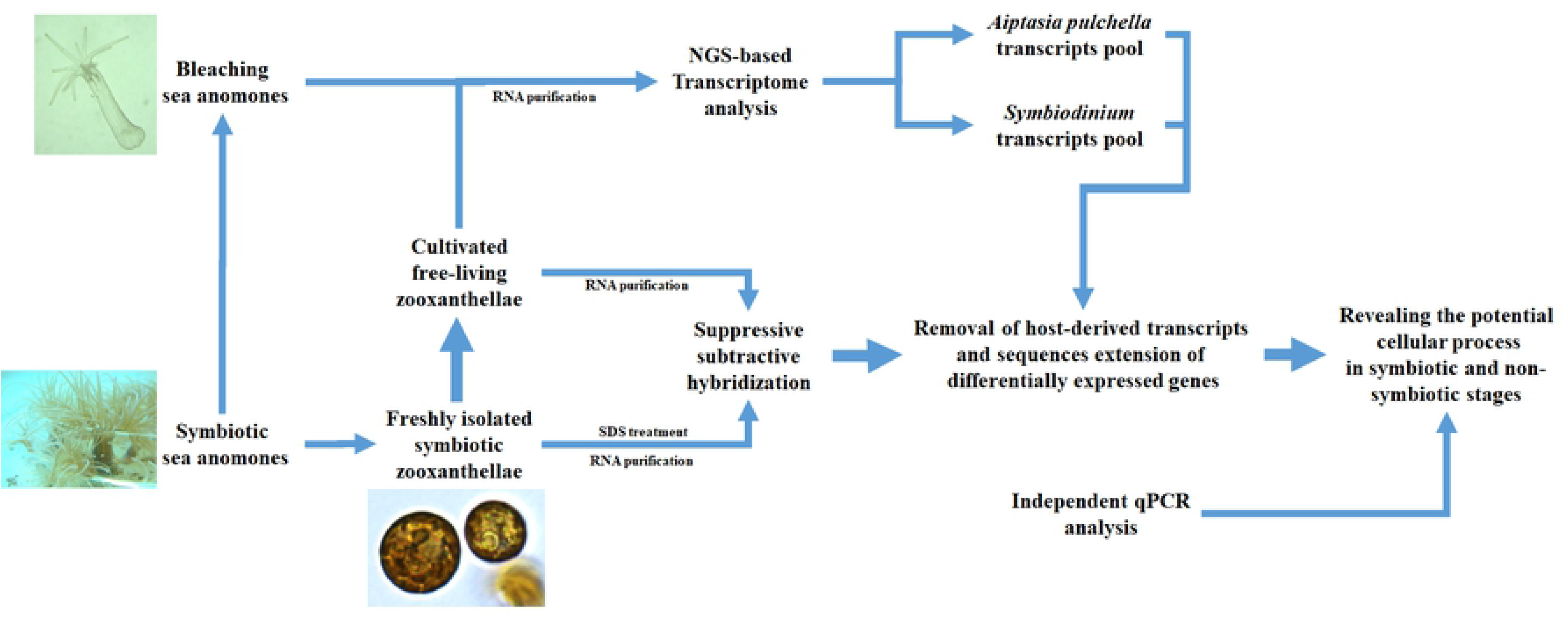
Schematic representation of SDS treatment, transcriptome sequencing, assembly, separation, and extension of host/symbiont sequences.

**Fig 3.**
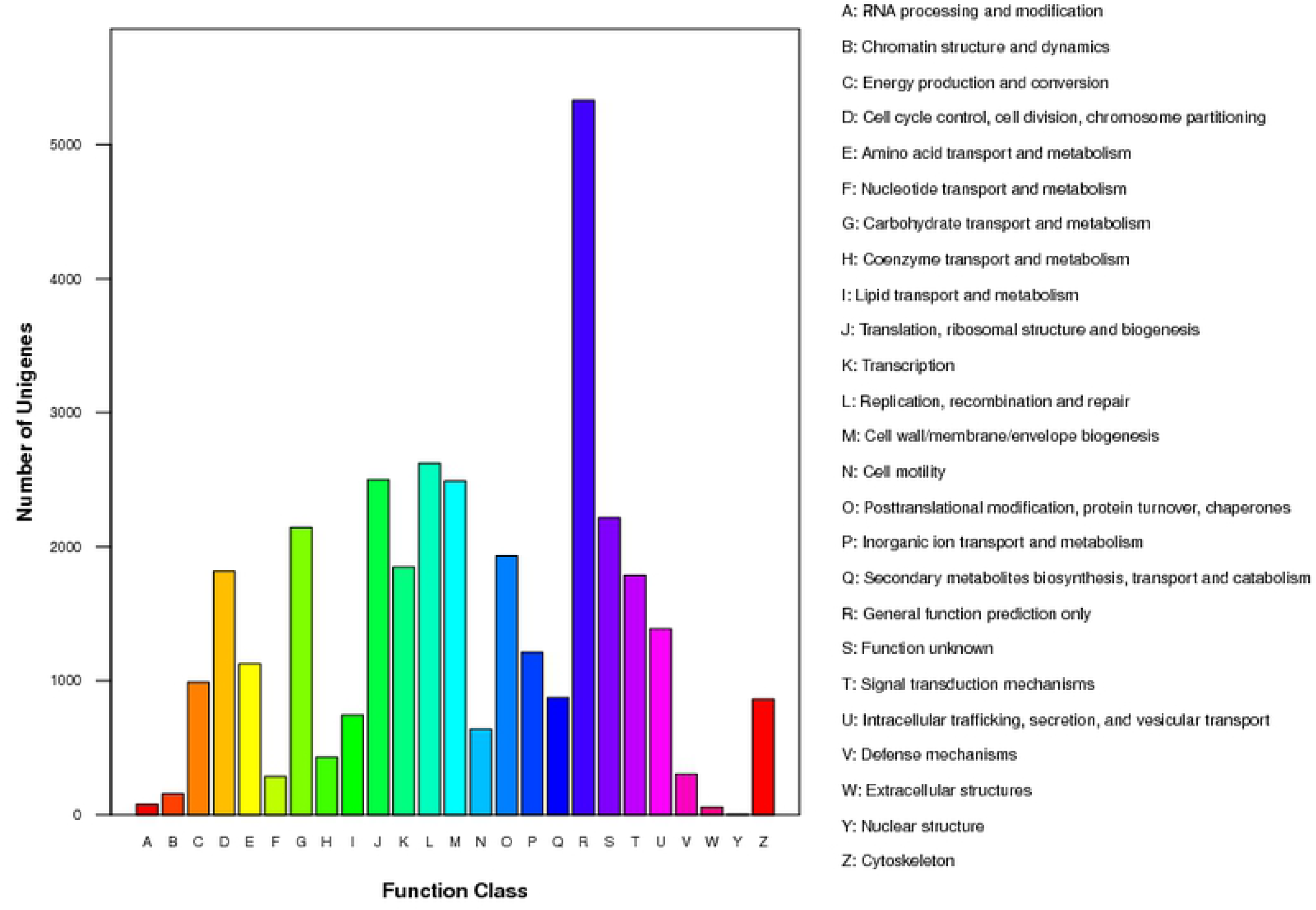
Clusters of Orthologous Genes classification of free-living *Symbiodinium* clade B expressing transcripts. The pathway contains: metabolism, genetic processing, environmental responses, cellular processes and organismal systems. Human diseases results were removed in the result of KOG pathways.

**Fig 4.**
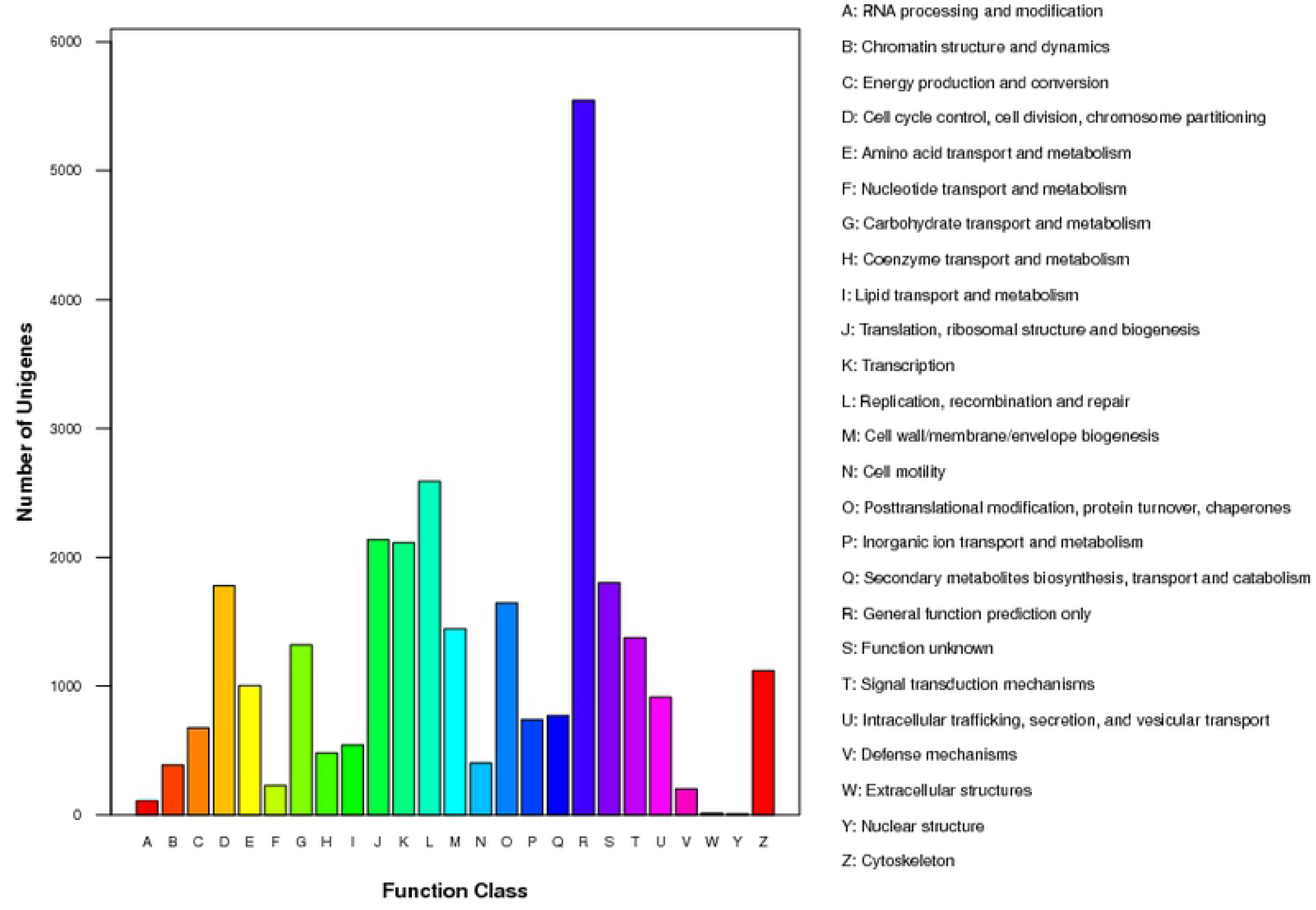
Cluster of Orthologous Genes classification of bleaching *Aiptasia pulchella* expressing transcripts. The pathway contains: metabolism, genetic processing, environmental responses, cellular processes and organismal systems. Human diseases results were removed in the result of KOG pathways.

**Fig 5.**
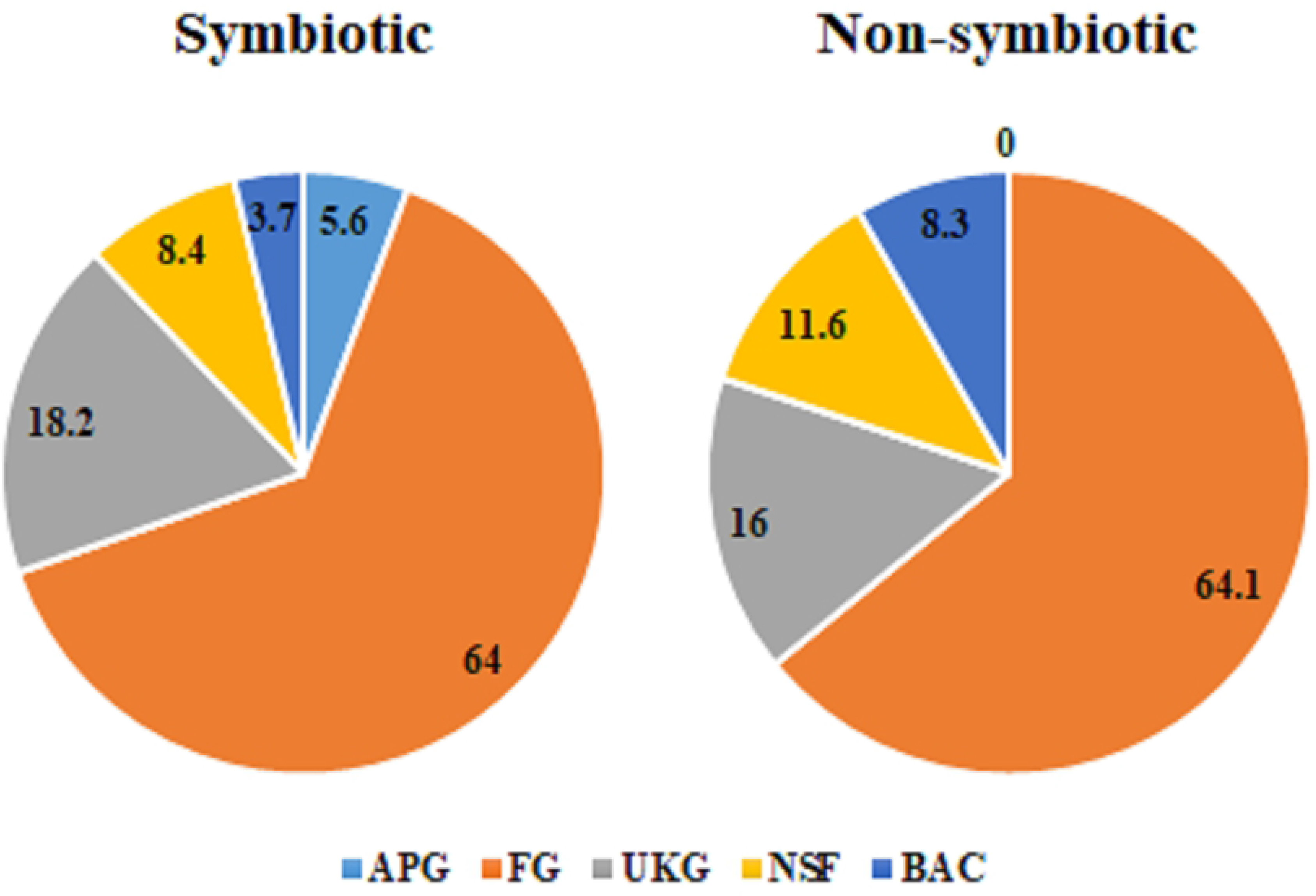
Taxonomy based on transcriptome data generated from *Aiptasia pulchella* - associated zooxanthellae. Taxonomies are provided from sequences of symbiotic and nonsymbiotic zooxanthellae through transcriptome analysis and suppressive subtractive hybridization. A strict and inclusive taxonomy is displayed. Screening revealed 214 symbiosis-specific expressed transcripts, displaying identity to only 5.6% of host-derived transcripts, 64% well-known functional genes. In the nonsymbiotic stage, the results indicated 181 differentially expressed transcripts, of which 64.1% belonged to well-known functional genes. ‘APG’ represents sequences of *Aiptasia pulchella* genes; ‘FG’ refers to genes with specific functions; ‘UKG’ represents genes of unknown function; ‘NSF’ indicates no similarity; ‘BAC’ refers to bacterial-like genes.

### Screening of differentially expressed transcripts between endosymbiosis

After SSH and sequence extension, the number of transcripts corresponding with specific functional genes was identified through BLAST with the NCBI GenBank database. Overall, 202 genes were differentially expressed in the symbiotic stage, of which 137 are genes had known functions, whereas the remaining 39 had unknown functions, 18 sequences did not display any similarity, and 8 genes were potentially bacterial genes. Otherwise, 181 genes were differentially expressed in the nonsymbiotic stage, of which 116 had known functions, 29 had unknown functions, 21 sequences did not share any similarity, and 15 genes were potentially bacterial genes (Fig 5). This study is focused on the 137 and 116 differentially expressed genes in the symbiotic and nonsymbiotic stages, respectively; they were divided into 7 important categories to elucidate the potential intracellular responses of endosymbiosis, as detailed below (Tables 1 and 2; detailed results shown in Tables S1 and S2).

**Table 1.**
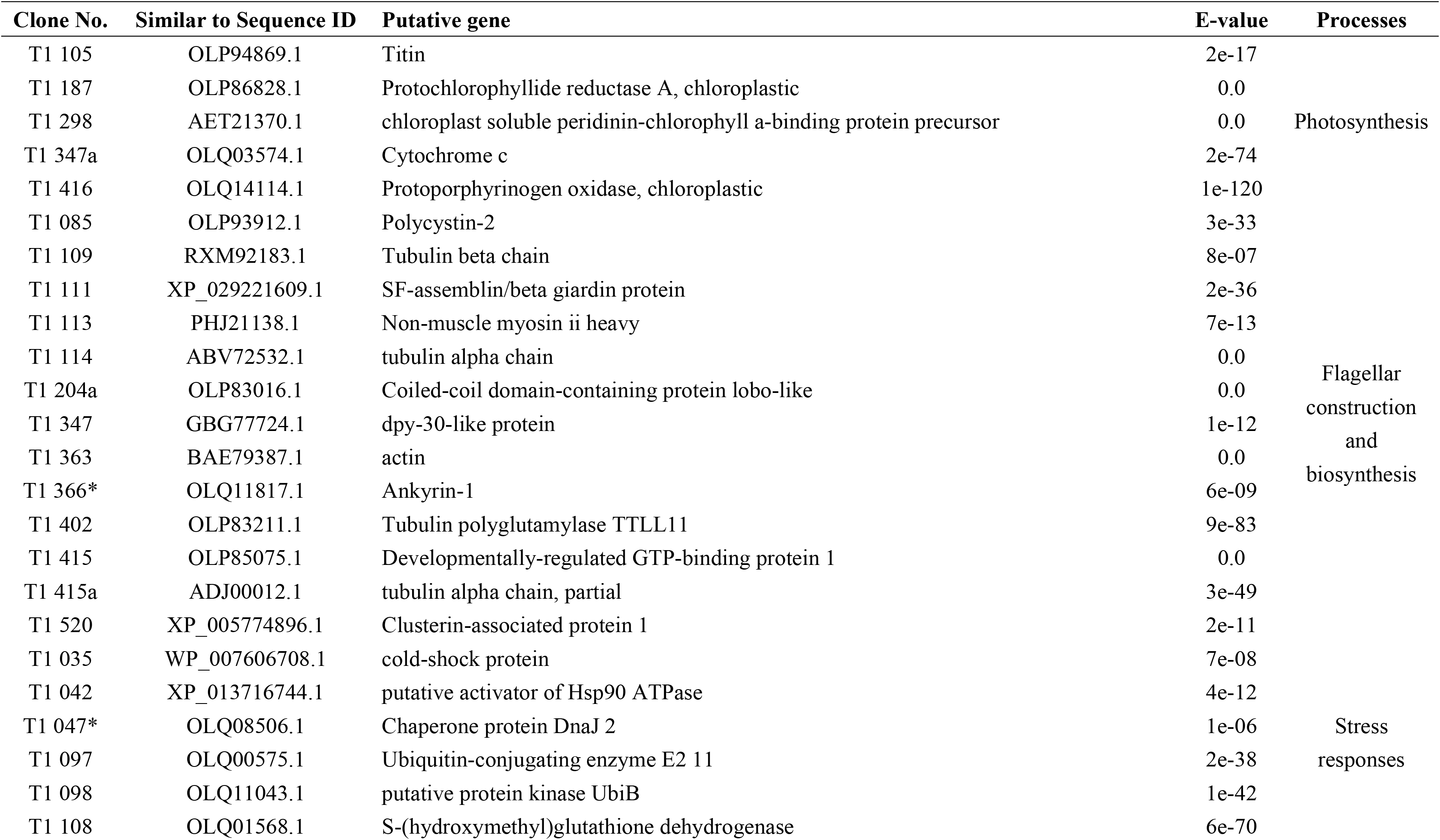
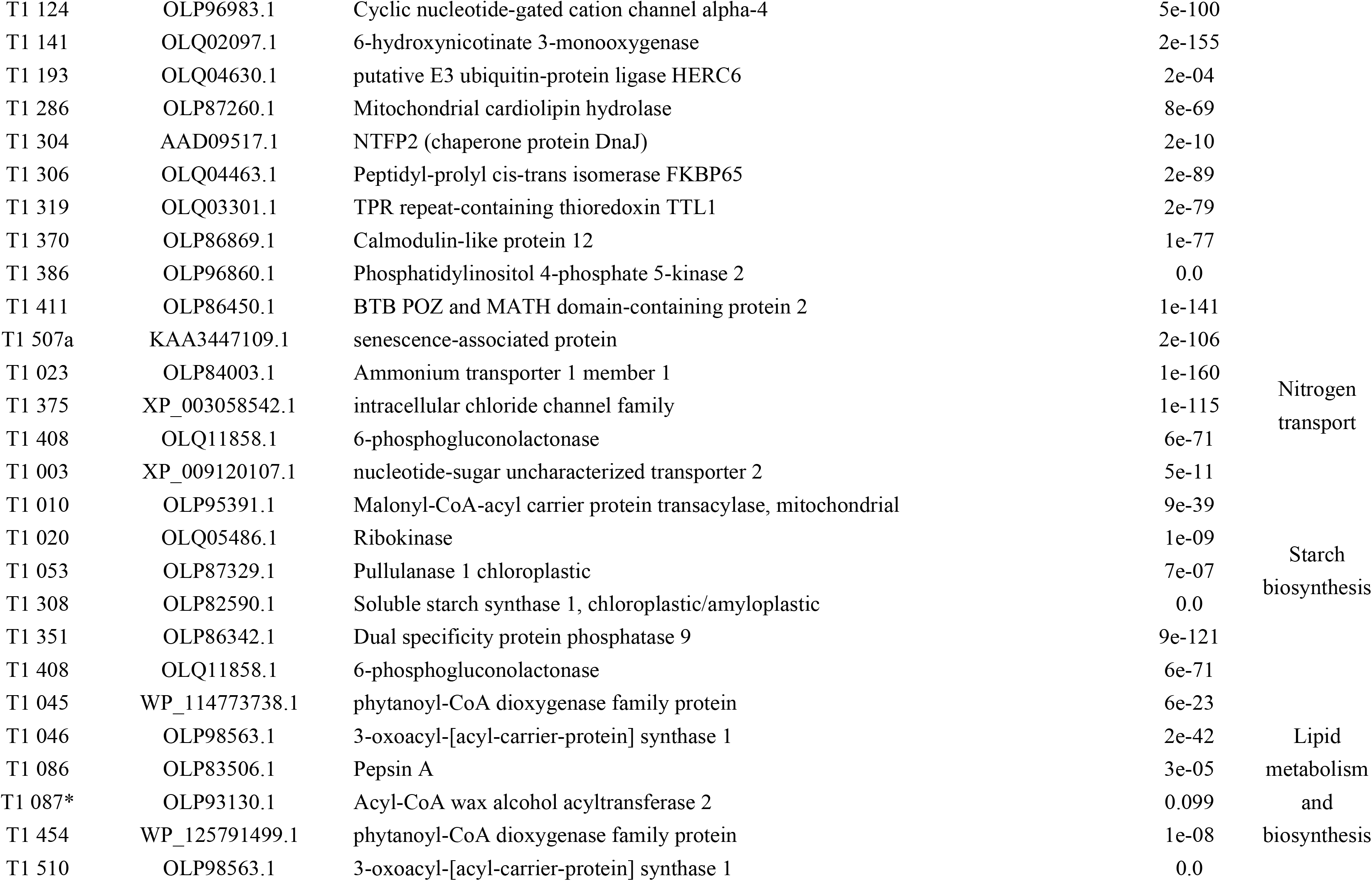
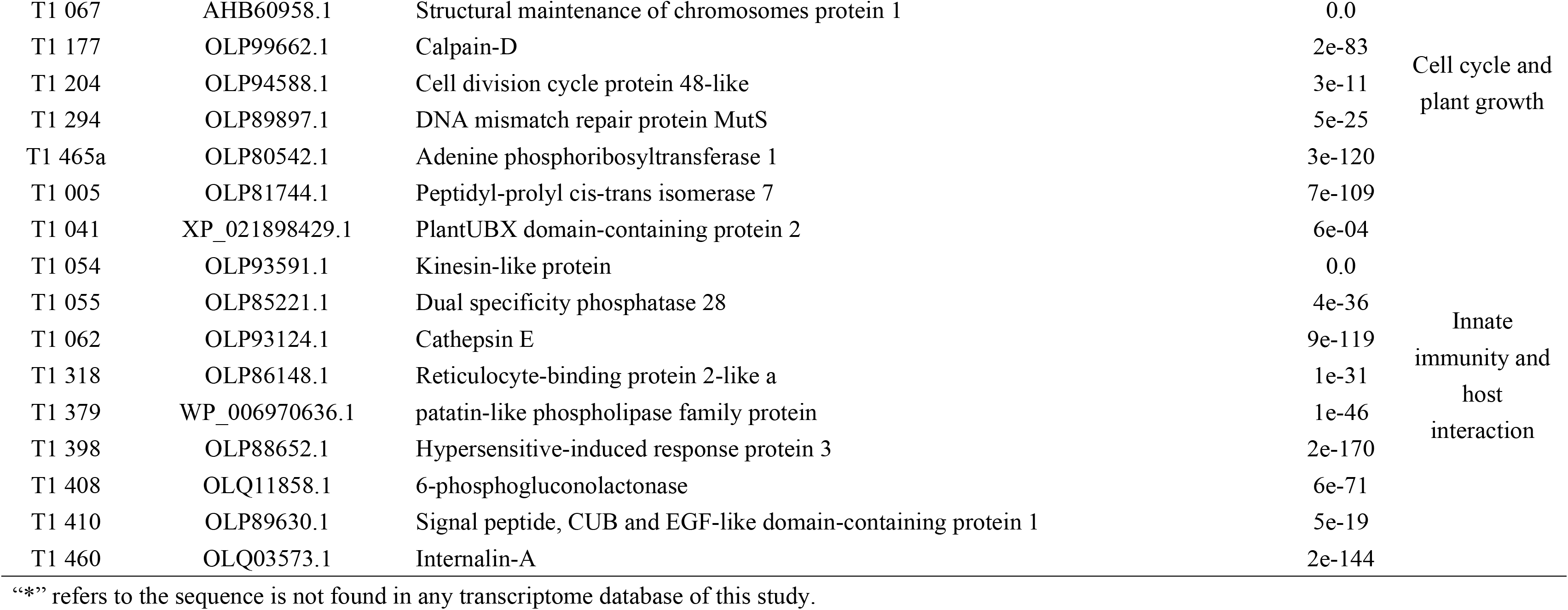
List of differentially expressed transcripts in symbiotic zooxanthellae.

**Table 2.**
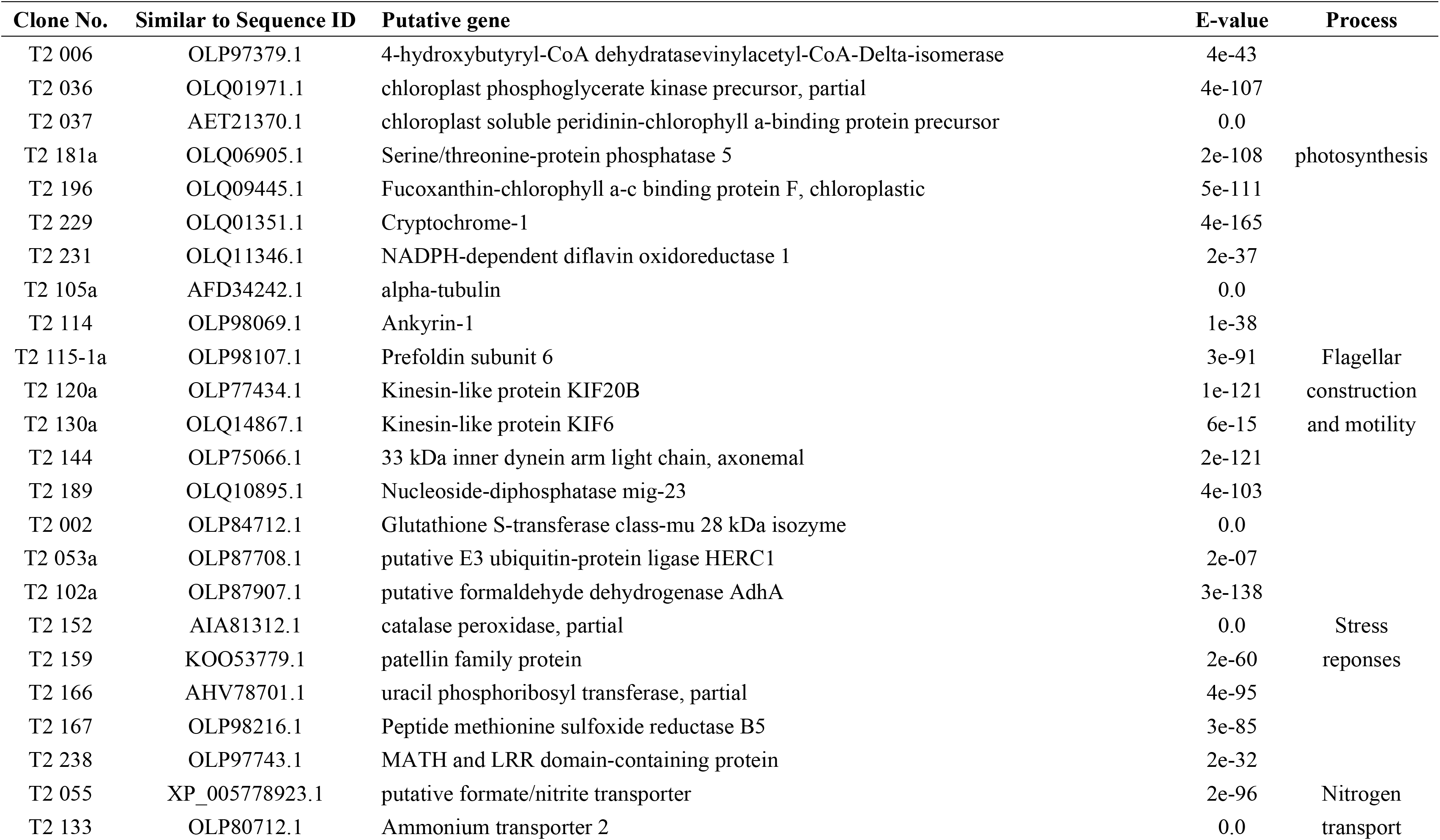
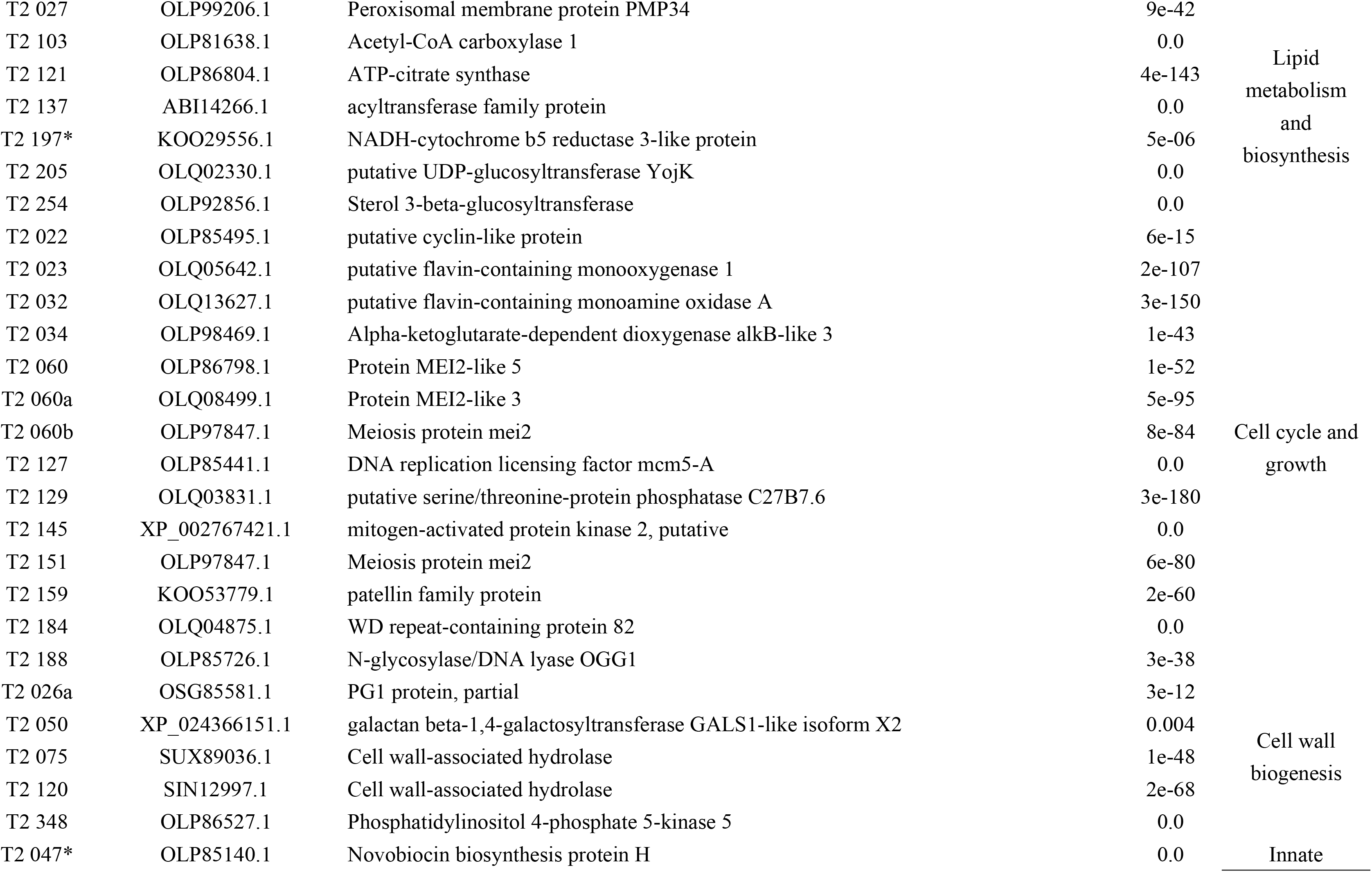
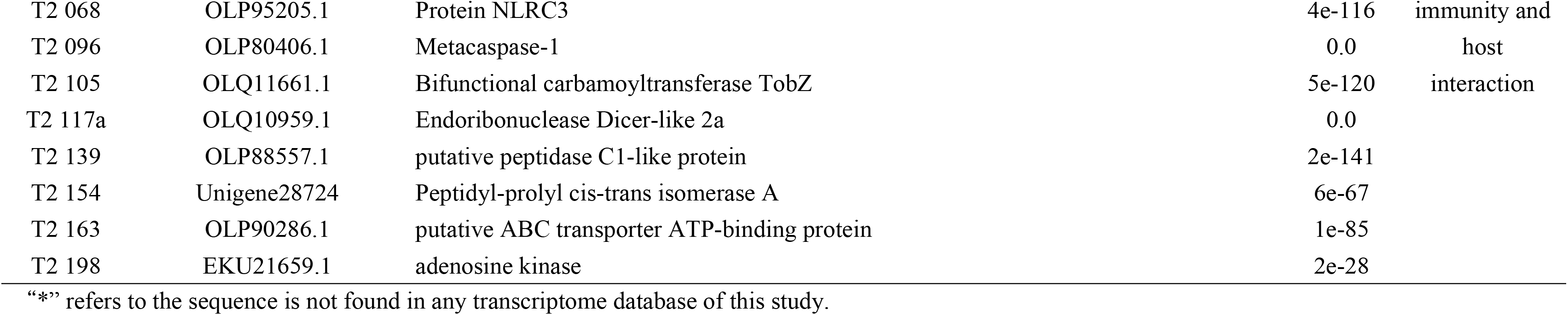
List of differentially expressed transcripts in nonsymbiotic zooxanthellae.

### Symbiotic zooxanthellae may produce chlorophyll to enhance photosynthesis

During photosynthesis, 5 differentially expressed genes detected in symbiotic zooxanthellae were implicated in chlorophyll biosynthesis. Table 1 shows that upregulated chloroplastic protoporphyrinogen oxidase eliminated protons from protoporphyrinogen IX to produce protoporphyrin IX, which was transformed into the pro-precursor of chlorophyll a, named protochlorophyllide (Pchlide) [53,54]. Further, light- and NADPH-dependent chloroplastic protochlorophyllide reductase A (4.8-time expression) promoted the phototransformation of protochlorophyllide to chlorophyllide (Chlide) [55], which are the biosynthetic precursors of chlorophyll a. Biosynthesis of chlorophyll a harvests solar energy from chloroplast soluble peridinin-chlorophyll a-binding protein and transport through cytochrome c-mediated electron transport [56,57], thus being essential for normal plant growth and development.

However, free-living zooxanthellae displayed different activities during photosynthesis. Expression of chloroplast phosphoglycerate kinase was implicated in the downregulation of photosynthetic activity during glycolysis impairment for metabolic adjustment and to optimize growth [58]. Chloroplastic fucoxanthin-chlorophyll a-c binding protein F, cryptochrome-1 and NADPH-dependent diflavin oxidoreductase 1 were differentially expressed (Table 2), suggesting that light responses of zooxanthellae were mediated by cryptochrome-1 [59], NADPH-dependent diflavin oxidoreductase 1 catalysed the NADP-dependent reduction of cytochrome c, but not cytochrome P450 *in vitro* [60] and energy was transferred from the carotenoid and chlorophyll C (or B) to chlorophyll A and the photosynthetic reaction centres where chloroplastic Fucoxanthin-chlorophyll a-c binding protein F is used to synthesize ATP and reduce solar power [61]. Altogether, the present results indicate that zooxanthellae in the symbiotic stage had an enhanced photosynthetic efficiency, which was not observed in the nonsymbiotic stage.

### Zooxanthella expressed numerous genes involved in flagellar motility and construction

In the nonsymbiotic stage, zooxanthellae expressed 7 transcripts corresponding to the genes involved in the dynamics of flagellar motility (Table 2) and regulated by Kinesin II-mediated anterograde intraflagellar transport (IFT) [62]. Kinesin-like protein KIF6 and KIF20B, the plus-end-directed motor enzyme required to complete cytokinesis and cellular motor activity, is directed toward the microtubule’s plus end [63,64]. As observed, two flagella assembled on free-living zooxanthellae through the regulation of kinesin-associated protein and IFT. Axonemal 33-kDa inner dynein arm light chain (expressed at 1.36-fold; Table 3) and prefoldin subunit 6 potentially contribute to flagellar movement and the stable assembly of a subset of inner dynein arms for the binding of these arms to the outer doublet microtubules of the axoneme to facilitate swimming [65,66]. Inversely, SSH revealed 13 unique transcripts of zooxanthellae in the symbiotic stage, which were involved in flagellar function and biosynthesis (Table 1). These results are concurrent with those of a previous study wherein RNAi-mediated polycystin-2 silencing revealed defects in flagella-dependent mating in *Chlamydomonas reinhardtii* [67]. Coiled-coil domain-containing protein lobo-like is the primary component of the nexin-dynein regulatory complex (N-DRC), which is essential for N-DRC integrity and regulates flagellar motility [68]. Dpy-30-like protein markedly contributed to flagellar radial spokes, which contribute to the regulation of dynein arm activity and thus the pattern of flagellar bending [69]. Clusterin-associated protein 1 was required for cilial biogenesis and functioned within the multiple intraflagellar transport complex B (IFT-B) [70]. Impairments of SF-assemblin-beta giardin protein in *Chlamydomonas* resulted in defects in flagellar assembly [71,72]. Developmentally regulated GTP binding protein 1 as a microtubule-binding protein can diffuse on microtubules, promote their polymerization, drive microtubule formation into bundles, and stabilize microtubules *in vitro* [73]. Consistently, qPCR revealed that relative expression of nonmuscle myosin II heavy chain, SF-assemblin-beta giardin protein and alpha tubulin were 8.01-, 1.86-, and 11.46-fold. Interestingly, Tubulin polyglutamylase TTLL11 was also detected herein, since the well-known Polyglutamase, which preferentially modifies alpha-tubulin was required for CCSAP localization to both spindle and ciliary microtubules [74]. These results suggest that zooxanthella contained in the host cells contributed to the generation of the flagella or cilia and on membrane trafficking.

**Table 3.**
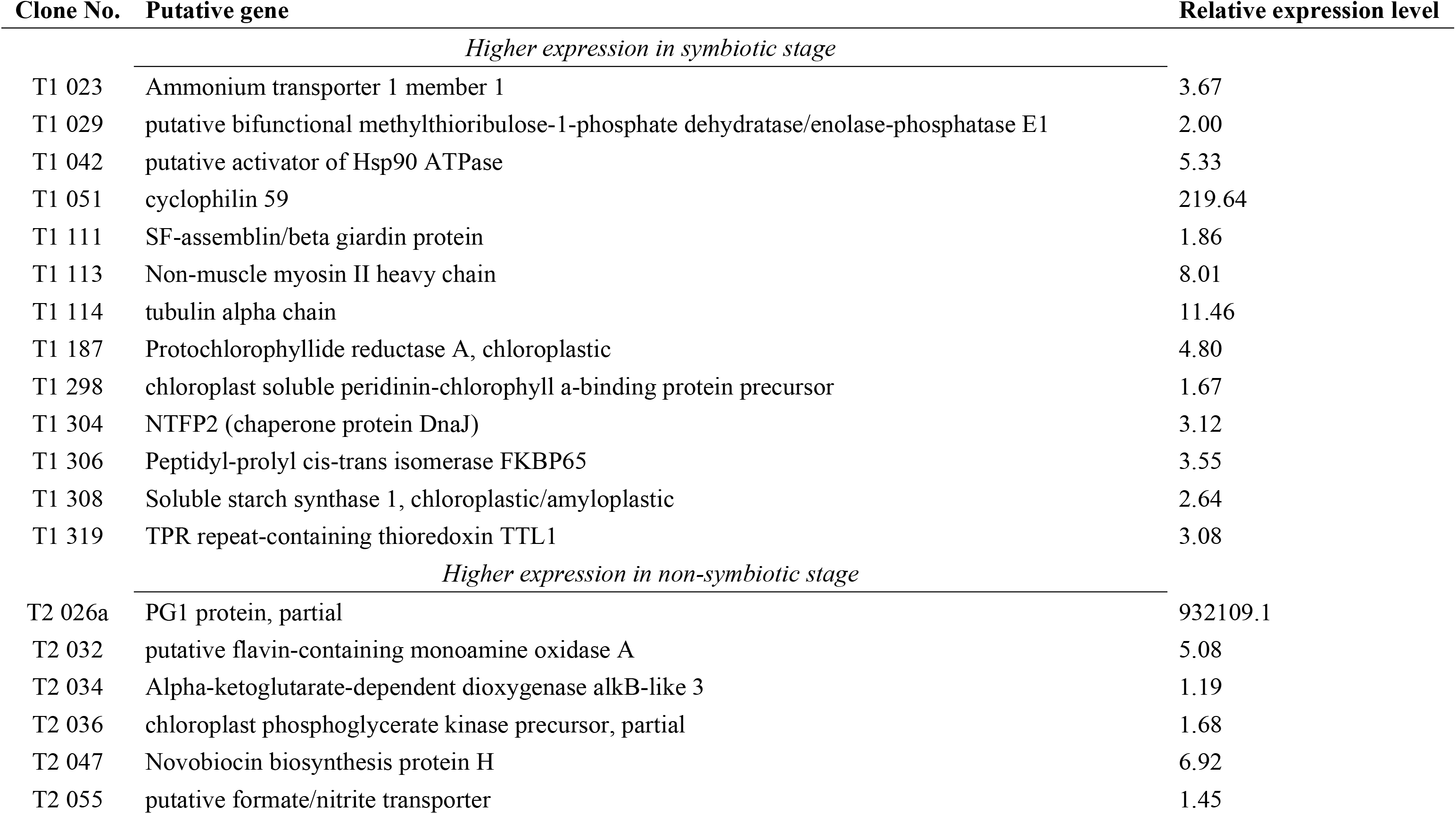
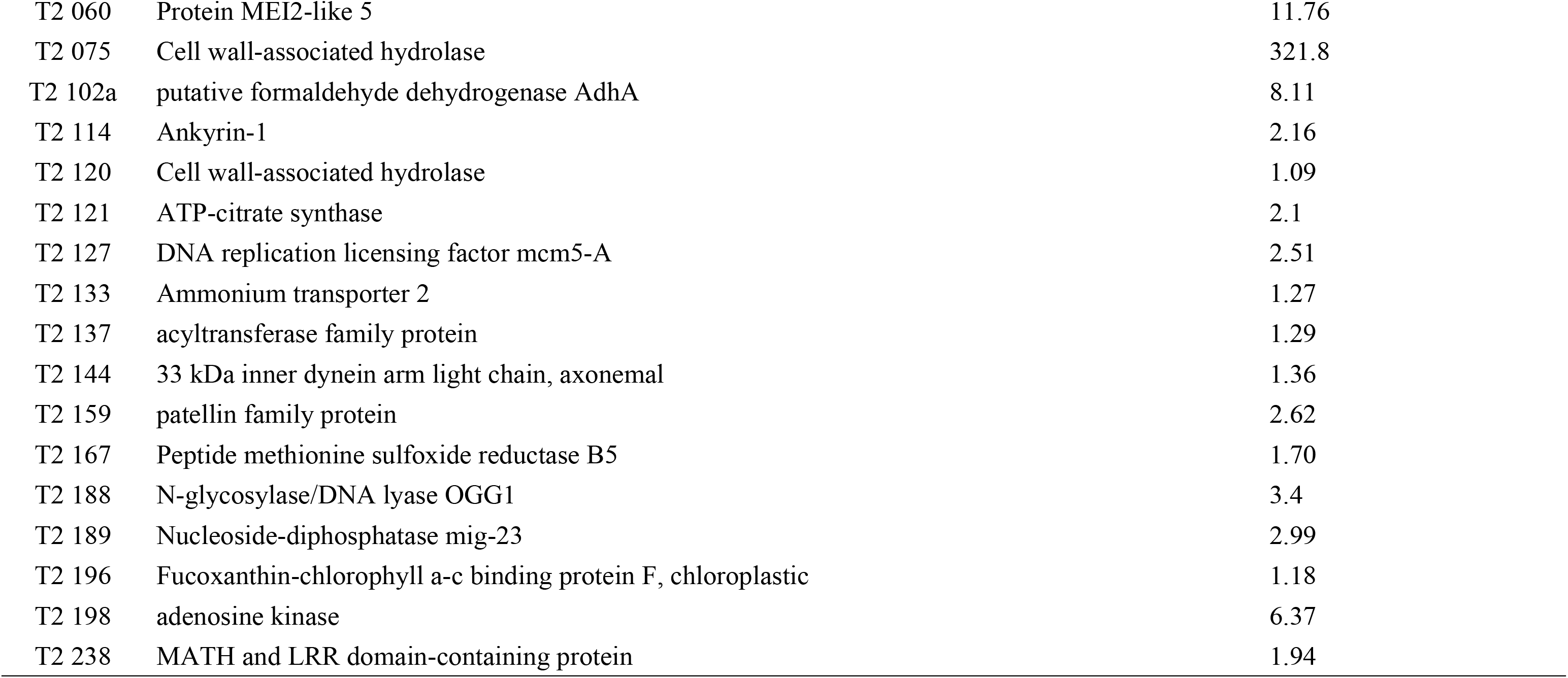
List of differentially expressed zooxanthellal genes revealed through quantitative PCR.

### Symbiotic zooxanthellae are potentially subjected to different stresses

BLAST analysis of 17 transcripts of zooxanthellae in the symbiotic stage corresponded to various stress-induced genes (Table 1), including cold-shock protein, putative activator of Hsp90 ATPase, NTFP2, Chaperone protein DnaJ 2, Ubiquitin-conjugating enzyme E2 11, putative protein kinase UbiB, S-(hydroxymethyl) glutathione dehydrogenase, Cyclic nucleotide-gated cation channel alpha-4, 6-hydroxynicotinate 3-monooxygenase, putative E3 ubiquitin-protein ligase HERC6, mitochondrial cardiolipin hydrolase, peptidyl-prolyl cis-trans isomerase FKBP65, TPR repeat-containing thioredoxin TTL1, calmodulin-like protein 12, phosphatidylinositol 4-phosphate 5-kinase 2, BTB POZ, MATH domain-containing protein 2, and senescence-associated protein. These genetic responses indicate that zooxanthellae are potentially under biotic and abiotic stress including senescence, cold and heat stress, oxidative stress, photodamage, nitrosative stress, salt stress, hyperosmotic stress, osmotic stress, and water stress. For example, expression of cold-shock protein, putative activator of Hsp90 ATPase, NTFP2, chaperone protein DnaJ 2, ubiquitin-conjugating enzyme E2 11, and putative E3 ubiquitin-protein ligase HERC6 are speculated to facilitate cell survival even at temperatures lower or higher than the optimum growth temperature [75–78]. Putative protein kinase UbiB can phosphorylate tocopherol cyclase VTE1 and help recycle oxidated alpha-tocopherol quinone to prevent chloroplast photo damage under continuous red light [79]. S-(hydroxymethyl) glutathione dehydrogenase modulates nitrosative stress during plant development [80]. Mitochondrial cardiolipin hydrolase has been speculated to directly contribute to the processing of primary piRNA transcripts to regulate mitochondria under hyperosmotic stress [81]. TPR repeat-containing thioredoxin TTL1 potentially serves as a positive regulator of ABA signalling during germination under osmotic stress [82]. Plant-type phosphatidylinositol-4-phosphate 5-kinases and BTB POZ, and MATH domain-containing protein 2 were responsive to environmental factors, thus playing an essential role in coordinating plant growth under water stress [83]. Oxidative stress induced senescence-associated protein to have a genetically based program leading to cell senescence and death [84]. Furthermore, qPCR confirmed zooxanthellal NTFP2 and putative activator of Hsp90 ATPase in the symbiotic stage were upregulated (5.33-fold and 3.12-fold) rather than in the nonsymbiotic stage. However, fewer stress-induced responses were observed through differentially expressed transcripts among nonsymbiotic zooxanthellal (Table 2), suggesting that nonsymbiotic zooxanthellae may encounter phosphate limitations [85], pH stress [86], oxidative stress [87], and damage from antibiotics, which are derived from culture conditions. Overall, the results showed that symbiotic zooxanthellae are not in a completely mutually beneficial environment as we expected. To survive in the host cell, zooxanthellae may have to face a complicated microenvironment different from that of seawater; thus, they are subjected to various adverse reactions.

### Starch, not lipids, may be the major product of carbohydrate biosynthesis in symbiotic zooxanthellae

The major product of photosynthesis is probably transferred to starch, and not lipids, during *A. pulchella* and *Symbiodinium* endosymbiosis. This speculation was based on the SSH findings herein, which revealed 7 differentially expressed transcripts of zooxanthellae in the symbiotic stage (Table 1), including nucleotide-sugar uncharacterized transporter 2, malonyl-CoA-acyl carrier protein transacylase, ribokinase, chloroplastic pullulanase 1, chloroplastic/amyloplastic soluble starch synthase 1 and dual-specificity protein phosphatase 9, and 6-phosphogluconolactonase. Furthermore, qPCR analysis revealed a 2.64-fold expression level of symbiotic zooxanthellal chloroplastic/amyloplastic soluble starch synthase 1, which primarily contributes to starch synthesis [88], compared with nonsymbiotic zooxanthellae. Furthermore, symbiotic zooxanthellae expressed dual-specificity protein phosphatase 9 and chloroplastic pullulanase 1, a starch-associated phosphoglucan phosphatase and enzyme, both of which were involved in starch biosynthesis, catabolism, and degradation [89,90]. Indeed, the expression of 6-phosphogluconolactonase is potentially correlated with the requirement of sugar-dependent nitrate assimilation [91]. These results indicate that starch and starch-derived products may be the major product of carbohydrate biosynthesis in symbiotic zooxanthellae.

Interestingly, lipid biosynthesis and transport were probably the most prominent metabolic activity among nonsymbiotic zooxanthellae, instead of starch biosynthesis. Table 2 lists several differentially expressed transcripts involved in lipid biosynthesis, transport, and metabolism, including sterol 3-beta-glucosyltransferase, which catalyses the synthesis of steryl glycosides (SGs) and acyl steryl glycosides (ASGs), which are the most abundant sterol derivatives in higher plants [92]. NADH-cytochrome b5 reductase 3-like protein and acyltransferase family protein are involved in fatty acid desaturation and elongation and cholesterol biosynthesis [93,94]. Acetyl-CoA carboxylase 1 and ATP-citrate synthase couple energy metabolism with fatty acids synthesis to support cell growth [95,96]. Altogether, these results indicate that the production of lipids and lipid derivatives is potentially involved in cell proliferation to increase the zooxanthellae population in the nonsymbiotic stage.

### Symbiotic zooxanthellae potentially utilise nitrates and ammonium first, but not nitrites

Nonionic NH_3_ can freely passively diffuse through biological membranes; however, it is retained within the symbiosome owing to its acidic microenvironment in the lumen [25–28]. The nitrogen transport demand between the symbiosome and zooxanthellae cells is reflected through the expression of ammonium transporter 1 and intracellular chloride channel family protein (Table1 and 3), both of which play essential roles in ammonium and nitrate transport and their associated molecular responses [97–99]. Interestingly, zooxanthellae in the nonsymbiotic stage differentially expressed ammonium transporter 1 and putative formate nitrite transporter (1.45-fold) rather than in the symbiotic stage (Tables 1 and 3). These results indicate that nitrate and ammonium are absorbed on priority by symbiotic zooxanthellae; nonsymbiotic zooxanthellae, nitrite, and ammonium.

### Nonsymbiotic zooxanthellae potentially mediate cell cycle progression and cell wall biosynthesis to accelerate cell growth

Only 6 transcripts of zooxanthellae were differentially expressed in the symbiotic stage (Table 1), and BLAST analysis revealed transcripts including structural maintenance of chromosomes protein 1 (also known as ‘cohesion complex subunit SMC-1’), calpain-D, calpain-15 isoform X2, PBS lyase cell division cycle protein 48-like, DNA mismatch repair protein MutS, and adenine phosphoribosyltransferase 1. The cohesin complex potentially forms a large proteinaceous ring within which sister chromatids can be trapped, playing an important role in spindle pole assembly during mitosis and in chromosome migration and regulating chromosome segregation and cell division during embryogenesis [100,101]. Calpain-D and calpain-15 isoform X2, both being plant phytocalpains, are key regulators of cell proliferation and differentiation during plant organogenesis, partly acting by regulating the CycD/Rb pathway [102]. PBS lyase cell division cycle protein 48-like protein probably functions in cell division, and adenine phosphoribosyltransferase 1 converts cytokinins from free bases to their corresponding nucleotides [103,104]. DNA mismatch repair protein MutS is involved in DNA mismatch repair. Obviously, the primary cell cycle activities in symbiotic zooxanthellae were potentially correlated with the maintenance of chromosome structure and differentiation, concurrent with cell cycle arrest of symbiotic zooxanthellae in the G_1_/G_0_ phase.

Compared with symbiotic zooxanthellae, nonsymbiotic zooxanthellae displayed activities associated with cell cycle transition, meiosis, initiation and elongation phases of DNA replication, cell proliferation and differentiation, cytokinesis, regulation of chromatin structure, and repair of damaged DNA. The transcripts responsible for cell cycle regulation were similar to those of putative cyclin-like protein, putative flavin-containing monooxygenase 1, alpha-ketoglutarate-dependent dioxygenase alkB-like 3, protein MEI2-like 5, protein MEI2-like 3, meiosis protein mei2, DNA replication licensing factor mcm5-A, putative serine threonine-protein phosphatase C27B7.6, mitogen-activated protein kinase 2, patellin family protein, and WD repeat-containing protein 82. As previously reported, putative cyclin-like protein and WD repeat-containing protein 82 are the potential primary regulators of cyclin-dependent kinases, playing key roles in regulating mitotic cell cycle transition [105,106]. DNA replication licensing factor mcm5-A is a putative replicative helicase essential for the initiation and elongation phases of DNA replication in plant cells [107]. Putative serine threonine-protein phosphatase C27B7.6 regulates phosphorylation involved in cell proliferation, programmed cell death, and cell differentiation [108]. Protein MEI2-like 5, protein MEI2-like 3, meiosis protein mei2, and mitogen-activated protein kinase 2 were potential RNA-binding proteins essential for commitment to meiosis [109,110]. The patellin family protein is a carrier protein potentially involved in membrane-trafficking events associated with cell plate formation during cytokinesis [111]. All differentially expressed genes in nonsymbiotic zooxanthellae increase cell division in the presence of sufficient nutrient supplements and space during culture.

Especially, five differentially expressed transcripts of nonsymbiotic zooxanthellae were implicated in cell wall biosynthesis, which is essential for cell division, thus increasing the zooxanthellae population. In particular, PG1 protein expression levels in nonsymbiotic zooxanthellae were 93,216-fold that in symbiotic zooxanthellae (Table 3). The PG1 protein belongs to polygalacturonases (PGs), which act on the homogalacturonan chain, thus softening the cell wall [112] and potentially promoting cell division in nonsymbiotic zooxanthellae during reproduction. Several genes involved in PG modification and pectin biosynthesis were detected (Table 2), such as a cell wall-associated hydrolase with a specific role in modifying PG [113]. Galactan beta-1,4-galactosyltransferase GALS1-like isoform X2 and phosphatidylinositol 4-phosphate 5-kinase 5 both regulated pectin biosynthesis and secretion [114,115]. Robust cell wall regulation in zooxanthellae may have occurred owing to frequent cell division to increase the population of nonsymbiotic zooxanthellae; however, this effect was not observed in symbiotic zooxanthellae, and no cell-wall-related transcripts were found by the SSH assay; this result is concurrent with cell cycle arrest in the G_1_/G_0_ transition [31].

### Symbiotic zooxanthellae expressing immune-related genes associated with host defence

Residing in the *Aiptasia* host may present challenges associated with the intracellular defence mechanism in host cells, as described previously [35–37]. As predicted, SSH screening revealed numerous transcripts corresponding to those involved in innate immunity, inflammation, virus–host interactions, and antifungal defence, including peptidyl-prolyl cis-trans isomerase 7 (alternatively named as cyclophilin-7), which potentially regulate the responses to inflammatory diseases including atherosclerosis and arthritis and viral infections [116]. Similar to cathepsin D, cathepsin E has slightly broader specificity and potentially contributes to immune responses [117]. Kinesin-like protein and patatin-like phospholipase family protein are involved in microbial infections with fatty acid metabolism and immune regulation by apicomplexan and plants [118,119]. Dual-specificity phosphatase 28 limits the spread of the HR response after infection by necrotrophic pathogens including *Botrytis cinereal* [120]. For fungus-plant interactions, UBX domain-containing protein 2 and hypersensitive-induced response protein 3 negatively regulate the powdery mildew-plant interaction [121] and contribute to plant basal resistance via an EDS1 and SA dependent pathway [122,123]. In particular, internalin A was differentially expressed in symbiotic zooxanthellae. As reported, internalin A mediates the entry of *L. monocytogenes* into host intestinal epithelial cells and internalin A facilitates *L. innocua* uptake by host cells [124,125]. Importantly, symbiotic zooxanthellae presented transcripts similar to gene groups associated with the immune response, suggesting that the communication between the host and symbionts is potentially associated with immune regulation, which is one of the key factors responsible for symbiosis.

Other transcripts corresponded with numerous functional genes and were distinguished into other cellular processes (Tables 1 and 2). They may provide further evidence regarding the potential response during *A. pulchella-Symbiodinium* endosymbiosis.

### Numerous genes of unknown function were differentially expressed between symbiotic and nonsymbiotic stages

During genetic screening, various genes of unknown function were expressed in both symbiotic and nonsymbiotic stages of zooxanthellae. In total, 57 symbiotic and 50 nonsymbiotic expressed genes with unknown function were identified, of which 18/21 sequences in symbiotic/nonsymbiotic zooxanthellae did not share any similarity with sequences in NCBI GenBank, implying the lack of information regarding the genes associated with endosymbiosis; furthermore, quantitative analysis revealed that these unknown genes even provide evidence of >10-fold expression; hence, the present results provide more information regarding the molecular regulation of endosymbiosis.

## Conclusion

SSH is a potent, large-scale method for identifying differentially expressed target genes in symbiotic/nonsymbiotic zooxanthellae. This study showed that when purifying symbiotic zooxanthellae harboured in host cells, washing with 0.02% SDS effectively reduces the proportion of host RNA contamination during RNA extraction (by approximately 5.6%, Fig 5) and does not significantly influence gene expression in zooxanthellal PCP, HSP90, and actin. Therefore, SDS does not excessively affect gene expression in symbiotic zooxanthellae. Furthermore, combinatorial screening with SSH and sequence extension based on the NGS-based zooxanthellal gene database, symbiotic zooxanthellae focused on chlorophyll production during symbiosis to enhance photosynthesis. The carbohydrates produced through photosynthesis primarily include starch transported into the host cells for use. Furthermore, the symbiosome microenvironment may be subjected to stress in zooxanthellae. Numerous genes are associated with stress responses and are actively expressed herein for protection. Furthermore, the genes and channel proteins involved in immune regulation are potentially primarily implicated in the communication between the host and symbionts to the establishment and maintenance of endosymbiosis, including the cell cycle, which is arrested in the G_1_/G_0_ stage. Inversely, when leaving the host to live freely in seawater, photosynthesis in nonsymbiotic zooxanthellae primarily depends on the electron transport chain for photosynthesis progression, and its products are primarily converted into lipid forms for their own metabolic utilization and storage. Furthermore, genes associated with flagellar movements are expressed, thus facilitating smooth swimming. Simultaneously, to increase the zooxanthellae population, the acceleration of the cell cycle increased the rate of reproduction and population growth and is correlated with upregulated proteins including pG1, which is involved in the cell wall biosynthesis. Altogether, the present results provide insights into the mechanisms underlying physiological processes and molecular responses between symbiotic and nonsymbiotic zooxanthellae.

## Acknowledgments

This work supported by Ministry of Science and Technology, Taiwan (MOST 107-2311-B-992-003-MY3 and MOST 106-2313-B-992-301-). The funders had no role in study design, data collection and analysis, decision to publish, or preparation of the manuscript. The manuscript was edited by Wallace Academic Editing.

## Supporting information

Table S1. List of all differentially expressed transcripts in symbiotic zooxanthellae.

Table S2. List of all differentially expressed transcripts in non-symbiotic zooxanthellae.

